# DIscBIO: a user-friendly pipeline for biomarker discovery in single-cell transcriptomics

**DOI:** 10.1101/700989

**Authors:** Salim Ghannoum, Waldir Leoncio Netto, Damiano Fantini, Benjamin Ragan-Kelley, Amirabbas Parizadeh, Emma Jonasson, Anders Ståhlberg, Hesso Farhan, Alvaro Köhn-Luque

**Affiliations:** Department of Molecular Medicine, Institute of Basic Medical Sciences, University of Oslo, Norway; Oslo Centre for Biostatistics and Epidemiology, Faculty of Medicine, University of Oslo, Norway; Department of Urology, Northwestern University, Chicago, USA; Simula Research Laboratory, Lysaker, Norway; Sahlgrenska Center of Cancer Research, Department of Laboratory Medicine, Institute of Biomedicine, Sahlgrenska Academy at University of Gothenburg, Gothenburg, Sweden; Wallenberg Centre for Molecular and Translational Medicine, University of Gothenburg, Sweden; Department of Clinical Genetics and Genomics, Sahlgrenska University Hospital, Gothenburg, Sweden

**Keywords:** Single cell sequencing, Normalization, Gene filtering, ERCC spike-ins, Biomarkers, DEGs, Decision trees, Network analysis, Jupyter notebook, Binder

## Abstract

The growing attention toward the benefits of single-cell RNA sequencing (scRNA-seq) is leading to a myriad of computational packages for the analysis of different aspects of scRNA-seq data. For researchers without advanced programing skills, it is very challenging to combine several packages in order to perform the desired analysis in a simple and reproducible way. Here we present DIscBIO, an open-source, multi-algorithmic pipeline for easy, efficient and reproducible analysis of cellular sub-populations at the trasncriptomic level. The pipeline integrates multiple scRNA-seq packages and allows biomarker discovery with decision trees and gene enrichment analysis in network context using single-cell sequencing read counts through clustering and differential analysis. DIscBIO is freely available as an R package. It can be run either in command-line mode or through a computational pipeline using Jupyter notebooks. We also provide a user-friendly, cloud version of the notebook for researchers with very limited programming skills. We showcase all pipeline features using two scRNA-seq datasets. The first dataset consists of circulating tumor cells from patients with breast cancer. The second one is a cell cycle regulation datatset in myxoid liposarcoma. All analyses are available as notebooks that integrate in a sequential narrative R code with explanatory text and output data and images. These notebooks can be used as tutorials for training purposes and will guide researchers to explore their scRNA-seq data.

## 1. Introduction

Single-cell RNA sequencing (scRNA-seq) is a powerful technology that has already shown great potential and the field is rapidly expanding [1,2]. In animals, scRNA-seq is providing a leap forward in resolving cellular diversity and giving unprecedented insight into gene expression changes during development, regeneration and disease [3–8]. Recently, scRNA-seq also started to flourish in plant research [9,10]. The growing attention toward single-cell transcriptomics calls for the development of computational tools to interactively analyze scRNA-seq data for any target organism. Multiple tools for scRNA-seq analysis are being developed [11,12]. Most of them are characterized by specific analytic scopes in the wide spectrum of scRNA-seq analysis [13], thus leading many researchers to integrate several tools to address their questions.

RaceID is an excellent tool developed by Grün and colleagues to perform different aspects of scRNA-seq data analysis, including preprocessing, sub-population identification by *k*-means clustering, outlier cells detection and assessing differences in gene expression across clusters [14,15]. On the other hand, TSCAN is a remarkable tool for in silico pseudotime ordering implemented over cell clusters resulted mainly from model-based clustering (Ji and Ji 2016). Both RaceID and TSCAN do not handle external RNA controls consortium (ERCC) spike-in which can be used for accounting for cell-to-cell technical differences [16,17]. Furthermore, they lack features for biomarker discovery, such as networking, decision trees and gene enrichment analysis. The biomarker discovery approach enables researchers to find molecular markers and gene expression signatures for better diagnostic or prognostic techniques. Moreover, it could guide therapeutic decision-making [18]. Thus, multiple research efforts have been directed towards biomarker discovery and a number of computational tools are available. For instance, STRING is an online resource for networking analysis with known and predicted protein-protein interactions and functional enrichment analysis [19]. EnrichR is a prominent tool developed in the Ma’ayan lab for inferring knowledge about an input gene set by comparing it to annotated gene sets from over 160 libraries [20]. Decision trees are transparent and interpretable predictive models but they require considerable programming skills [21]. The RWeka package is a collection of machine learning algorithms, including decision tree analysis [22]. The rpart package is widely used for generating recursive partitioning trees [23]. For researchers with limited to no programming skills it can be very challenging to combine several of these tools to be able to perform a complete analysis. This is especially true for the designing of a complex computational pipeline that goes from single-cell sequencing read counts to biomarker discovery. Moreover, due to the required concatenation of heterogeneous programs and custom scripts via file-based inputs and outputs, as well as the program dependencies and version requirements, such a pipeline may suffer from reproducibility problems [24,25].

Hence, we developed DIscBIO (DIscovery of single-cell transcriptomics BIOmarkers). DIscBIO is an open-source, multi-algorithmic pipeline that provides an opportunity to analyze scRNA-seq data for any organism with a taxonomy ID. It allows biomarker discovery using decision trees and gene enrichment analysis in a network context from single-cell sequencing read counts through clustering and differential analysis. DIscBIO is implemented as an R package [26] published on the Comprehensive R Archive Network (CRAN). It can be run either in command-line mode or through a computational pipeline using Jupyter notebooks [27]. DIscBIO allows users to conveniently construct, analyze, visualize and tune scRNA-seq data interactively. DIscBIO notebooks integrate R code with explanatory text and output data and images in a sequential narrative. Our software allows users with limited programming skills to customize the pipeline in a stepwise fashion by tuning individual parameters, while experienced R programmers can fully edit and extend the pipeline. We also provide a cloud version using Binder [28] that allows users without programming skills to run the pipeline even without installing the programming language R, Jupyter, or any of the software that the pipeline integrates. In order to produce the cloud-hosted Binder version, the repository includes a specification of all the software used, which enables automatic or manual reproduction or modification of the necessary computational environment needed to execute the pipeline on any computational resources [28]. DIscBIO facilitates the creation of publication-ready plots for researchers with all levels of programming proficiency.

DIscBIO is open source, and it is freely available at https://github.com/ocbe-uio/DIscBIO. Its R package can be downloaded from https://cran.r-project.org/package=DIscBIO. To fully harness the potential of transcriptome analysis in deciphering complex diseases at single cell level, we showcase DIscBIO over two scRNA-seq datasets. The first dataset consists of circulating tumor cells (CTCs) from patients with breast cancer, the second one is from a myxoid liposarcoma cell line. In the first case, we investigate the connection between Golgi apparatus and oncogenesis of breast cancer to demonstrate how our analysis pipeline can be used to study this link based on current knowledge. In the second case, we focus on defining the molecular signature of sub-populations of myxoid liposarcoma cells in association with their cell cycle phases. Additionally, we showcase the analysis of a mouse scRNA-seq dataset with accession number GSE41265 [29] to demonstrate how to load and analyze datasets from the conquer repository [11]. The conquer repository provides access to analysis-ready scRNA-seq datasets from different species, including humans, mice and zebrafish.

## Pipeline description

DIscBIO is a multi-algorithmic pipeline for an easy, fast and efficient analysis of sub-populations and the molecular signatures that characterize them. The pipeline consists of four successive steps: 1) data pre-processing; 2) cellular clustering and pseudo-temporal ordering; 3) determining differentially expressed genes (DEGs); and 4) biomarker identification, including decision trees, networking and gene enrichment analysis (Figure 1). Below we summarize the sections and algorithms used in the pipeline. Detailed information may also be found in each of the sections of the notebooks:

**Figure 1.**
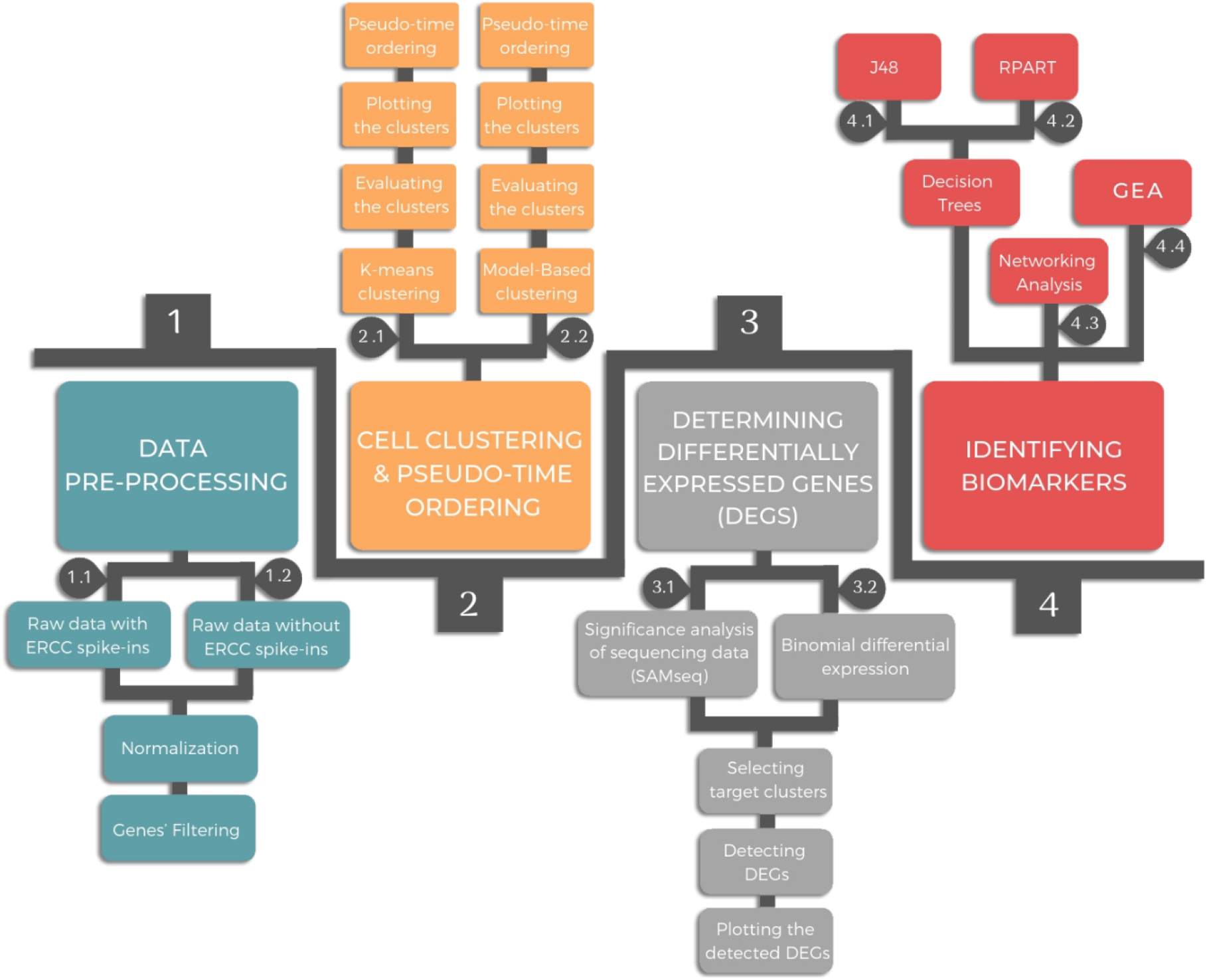
Overview of DIscBIO pipeline showing its four successive sections

### 1. Data pre-processing

Prior to applying data analysis, normalization and gene filtering are used to pre-process the raw read counts resulted from the sequencing. To account for RNA composition and sequencing depth among samples (single cells), the normalization method “median of ratios” is used. This method makes it possible to compare the normalized counts for each gene equally between samples. The key idea of gene filtering is to highlight the genes that manifest high variation across samples. DIscBIO provides three gene filtering techniques; noise filtering, expression filtering and designed filtering. In case the data include ERCC spike-ins, genes can be filtered based on variability in comparison to a noise level estimated from the ERCCs using an algorithm developed by Brennecke et al. [16]. Datasets without ERCCs, genes can be filtered either based on minimum expression in certain number of cells or based on a particular gene list of interest, for instance the genes involved in a desired cellular process.

### 2. Cellular clustering and pseudo-temporal ordering

Cellular clustering is performed according to gene expression profiles to detect cellular sub-populations with unique features. DIscBIO allows *k*-means clustering [30] using RaceID algorithm and model-based clustering [31] using TSCAN software. This pipeline estimates the number of clusters by finding the minimal clusters’ number at the saturation level of gap statistics, which standardizes the within-cluster dispersion [32]. DIscBIO enables robustness assessment of the detected clusters in terms of stability and consistency using Jaccard’s similarity statistics and silhouette coefficients [33,34]. To visualize the detected clusters, two common dimensionality reduction tools are implemented, principal component analysis (PCA) and *t*-distributed stochastic neighbor embedding (tSNE). DIscBIO uses the “tsne” package through RaceID to plot the clusters in tSNE maps [35,36]. Additionally, DIscBIO can detect outlier cells. The outlier identification is implemented based on RaceID using a background model based on distribution of transcript counts within a cluster. The background model is computed using the mean and the variance of the expression of each gene in a cluster. Outliers are defined as cells with a minimum number of outlier genes. This number is set by default to 2. In the case studies we set it to be 5% of the number of genes in the filtered dataset, this is based on the recommendation of De Vienne et al. [37]. Outlier genes are inferred from non-normalized transcript counts. Finally, pseudo-temporal ordering is implemented over the clusters resulted from the *k*-means and model-based clustering using TSCAN software. The pseudo-temporal ordering gradually orders cells based on their transcriptional profile, for example indicating the cellular differentiation degree.

### 3. Determining DEGs

Differences in gene expression between clusters are identified using significance analysis of sequencing data (SAMseq) [38] from the samr package [39]. SAMseq is a non-parametric statistical function dependent on Wilcoxon rank statistic that equalizes the sizes of the library by a resampling method accounting for the various sequencing depths. The analysis is implemented over the pure raw dataset that has the unnormalized expression read counts after excluding the ERCCs. The SAMseq function indirectly calls a subfunction called rankcol, which was written in Fortran. This function is used to attribute column-wise ranks to a matrix. Due to some limitations handling large datasets, the function was rewritten in R and incorporated into DIscBIO. Furthermore, binomial counting statistics are used to identify differentially expressed markers for each cluster. This is done by highlighting DEGs in the target cluster comparing to all the remaining clusters using binomial differential expression. Differentially expressed markers help characterize the molecular signatures. For instance, when analyzing cancer cells, it might help to characterize tumorigenic capabilities of each cell cluster leading to the development of better therapeutics [40]. The binomial differential expression is computed using RaceID through DESeq2 [41]. Volcano plots are used to visualize the results of differential expression analyses.

### 4. Identifying biomarkers

DIscBIO uses decision trees and hub detection through networking analysis to discover biomarkers. Decision trees are a very efficient classification technique in biomarkers discovery. In the current version of DIscBIO, two different decision trees, J48 and RPART, can be implemented to predict the sub-population of a target cell based on transcriptomic data. The J48 tree is implemented through the RWeka package whereas the RPART tree is computed using the rpart package. The performance of the generated trees can be evaluated for error estimation by ten-fold cross validation assessment. To identify protein-protein interactions we use STRING through its application programming interface. The outcome is used to test the connectivity degree and the betweenness centrality of the interaction network, which reflects the communication flow in the networks. Moreover, to gain mechanistic insight into DEGs gene and pathway enrichment analysis can be performed in DIscBIO through the enrichR package. Further information on how to use DIscBIO, possible options and documentation of the algorithms are included within our Jupyter notebooks.

## Case studies

To showcase DIscBIO, we analyzed two scRNA-seq datasets. One consisting of circulating tumor cells (CTCs) from patients with breast cancer, a second dataset from a myxoid liposarcoma cell line. Additionally, we illustrate how to use the conquer repository by loading and analyzing one of the available mice scRNA-seq dataset. For each case study, we provide a Jupyter notebook that includes the complete analysis and can be used as a guide to examine other datasets.

### 1. CTC case study

Here we analyze a dataset consisting of single (not clustered) CTCs collected from patients with breast cancer. Data are available in the GEO database with accession number GSE51827 [42]. The raw data includes RNA-seq from 29 samples, including 15 pools of single CTCs and 14 CTC-clusters that were obtained from blood of 10 breast cancer patients. Here, CTC-clusters were excluded and a dataset of 1462 single CTCs was used in the analysis. The CTC notebook is available at https://nbviewer.jupyter.org/github/ocbe-uio/DIscBIO/blob/dev/notebook/DIscBIO-CTCs-Notebook.ipynb. CTCs are a rare subset of cells found in the blood of cancer patients and arise as a consequence of tumors shedding cancer cells into blood vessels. They preserve primary tumor heterogeneity and imitate tumor characteristics [43]. CTCs may serve as metastatic seeds and succeed initiating secondary tumors. An increasing number of studies on CTCs focus on efforts to uncover and characterize molecular features of CTCs that predict their metastasis-generating potential. A prominent example is epithelial-mesenchymal transition [44]. Due to the widely accepted role of CTCs in metastasis, they can be used to showcase and explore potential involvement of molecular or cellular alterations in cancer progression. In our current case study, we aimed at 1) characterizing sub-populations of CTCs 2) linking alterations of Golgi apparatus with cancer progression to showcase how our analysis pipeline can be used to study this link based on current knowledge.

#### 1.1 Characterization of CTCs subpopulations

To characterize sub-populations of CTCs, the dataset was filtered based on the median expression of all genes in the dataset. Genes with less expression than the overall median expression in at least 10% of the cells were excluded. Cells with less than 1000 read counts were removed. In total, 215 genes and 1448 cells were used for further analysis. After filtering and computing gap statistics, four clusters were obtained based on *k*-means clustering (Figure 2a). Clusters 2 and 4 were stable and consistent. Clusters 1 and 3, instead, had low degree of stability and consistency. Pseudo-time ordering showed a clear separation between clusters 1 and 4 (Figure 2b). Likewise, the heatmap portrayal of cell-to-cell distances, where cluster centers were ordered by hierarchic clustering (Figure 2c), also exhibited a clear separation between clusters 1 and 4.

**Figure 2.**
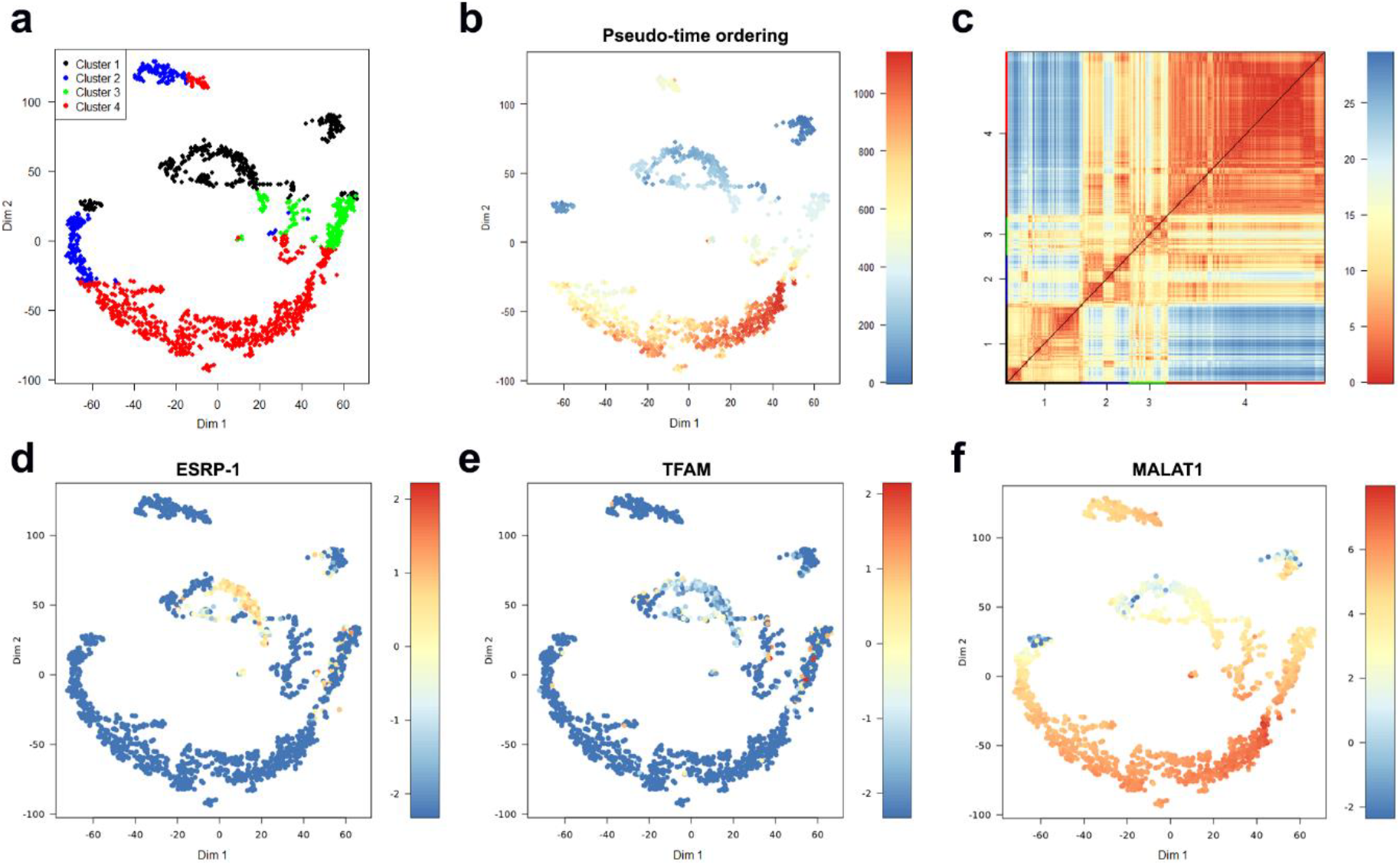
Identification of CTC sub-populations clustered based on the median expression of all genes in at least 10% of the cells. (a) *t*SNE map visualizing clusters of <u>circulating breast cancer cells</u> identified with *k*-means clustering. (b) The plot from (a) with cells colored based on their pseudo-time ordering. (c) A heatmap portrayal of cell-to-cell distances, cluster centers were ordered by hierarchic clustering. (d) The plot from (a) with cells colored based on their *ESRP-1* expression. (e) The plot from (a) with cells colored based on their *TFAM* expression. (f) The plot from (a) with cells colored based on their *MALAT1* expression.

Guha et al. (2018) have reported that metastasizing cells from a highly aggressive tumor exhibit higher frequency of mitochondrial defects. To investigate the mitochondrial defects across the clusters, we plotted the expressions of two genes reported by Guha et al: epithelial splicing regulatory protein (*ESRP*)-1 and mitochondrial transcription factor A (*TFAM*). Both genes were downregulated in cluster 4 compared to their expression in cluster 1 (Figure 2d-2e). Differentially expressed genes (DEGs) were extracted by two class unpaired response test using SAMseq, with significantly false discovery rate (FDR) less than 0.05. The gene enrichment analysis and the networking analysis performed over the list of the up-regulated DEGs in cluster 4 showed that many of these genes are involved in processes of relevance to cancer progression, such as exocytosis regulation, Rap1 signaling, and regulation of cell migration.

The networking analysis suggested five genes; *PF4, PPBP, ITGA2B, FERMT3* and *SELP* as central hub nodes (Figure 3a). The platelet factor 4 (*PF4*) is an endocrine factor with overexpression, associated with low survival of patients with lung cancer [45]. The Pro-Platelet Basic Protein (*PPBP*) is a member of the CXC subfamily of chemokines. *PPBP* was reported as an enhancer of the invasive ability of breast cancer cells [46]. Integrin alpha 2b (*ITGA2B*) is a member of the integrin family that regulates a diverse set of cellular processes crucial to the initiation, progression and metastasis of solid tumors, including breast cancer. The over-expression of *ITGA2B* promotes proliferation and invasion in breast cancer [47]. The *FERMT3* gene encodes a protein involved in integrin activation. *FERMT3* is known to enhance breast cancer progression and metastasis [48]. Selectin-P (*SELP*) was found to be important in organ-specific metastatic dissemination of breast cancer [49].

**Figure 3.**
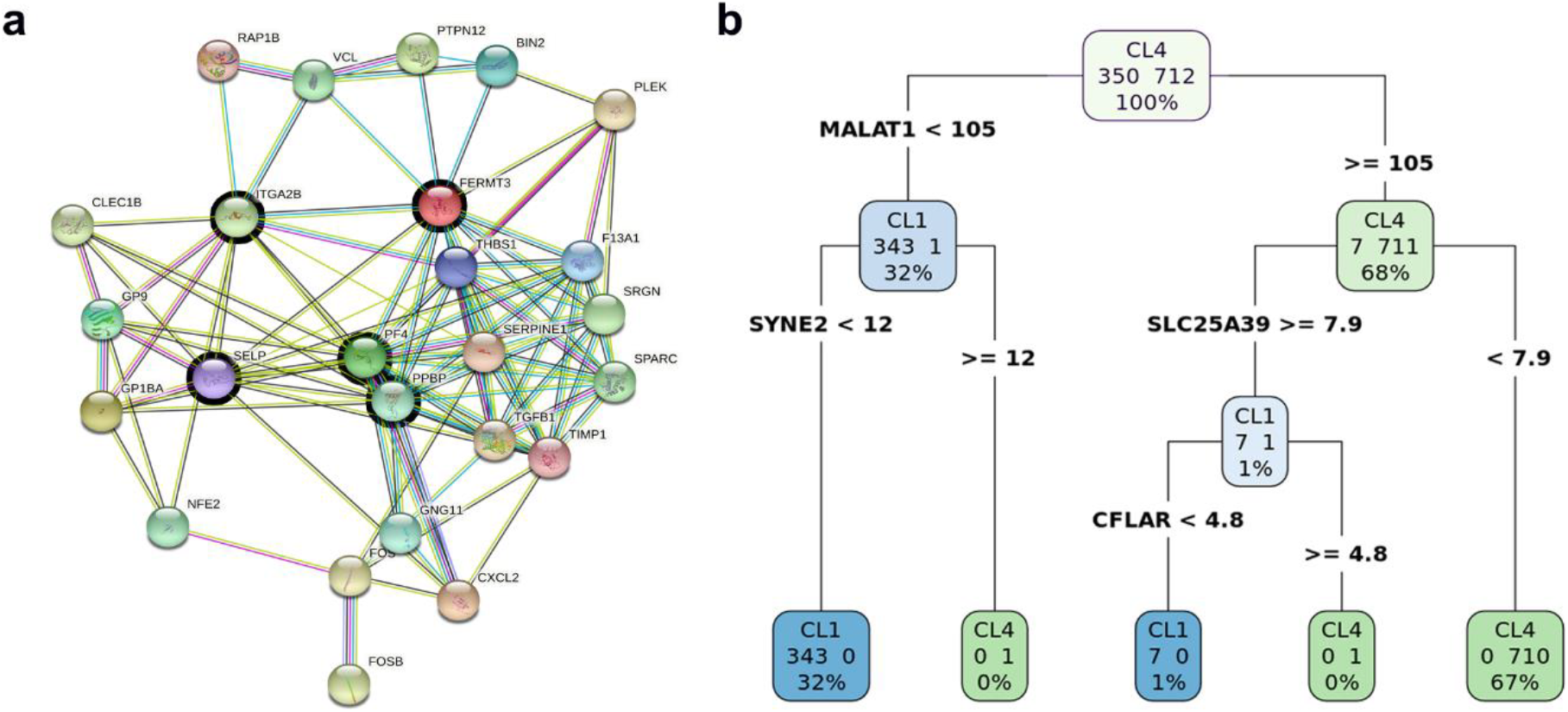
Biomarker discovery in cluster 4. (a) A sub-network of up-regulated DEGs in cluster 4. (b) A schematic figure explaining the RPART decision tree to predict the cluster identity of a target cell whether it belongs to cluster 1 or 4.

Moreover, binomial differential expression was performed to detect differentially expressed markers for each cluster. The metastasis associated lung adenocarcinoma transcript-1 (*MALAT1*) was detected to be a marker for cluster 4. *MALAT1* has been described as a long non-coding gene with contradictory functionality. Previous studies demonstrated that *MALAT1* promotes cell proliferation, migration, tumor growth, metastasis, and chemoresistance. On the other hand, a recent study by Kim and colleagues showed that *MALAT1* levels inversely correlate with breast cancer progression and metastatic ability in transgenic, xenograft, and syngeneic models [50]. All the CTCs in the selected dataset showed up-regulation of *MALAT1*, especially in cluster 4 (Figure 2f). Further investigation is needed to confirm the role of *MALAT1* in circulating breast cancer cells. The overall impact of our findings reported here is that cells in cluster 4 seem to have highly aggressive characteristics of invasion and metastasis comparing to less aggressiveness in cluster 1. Decision tree analysis was performed using the total list of the binomial DEGs to identify potential genes to predict the cluster identity of a target cell whether it belongs to cluster 1 or 4. The generated RPART decision tree (Figure 3b) included four decision nodes: *MALAT1, SYNE2, SLC25A39* and *CFLAR*. The performance metric of the RPART decision tree is remarkably high, with an accuracy of 99%, specificity of 100% and sensitivity of 99%.

#### 1.2 Linking alterations of Golgi apparatus with cancer progression

The Golgi apparatus is an evolutionary-conserved cellular organelle that plays a key role in protein sorting and post-translational modification, as well as being a hub for signaling molecules and to thereby contribute to the outcome of signaling cascades [51]. In addition, the Golgi has a role in regulating cell migration and cell polarity [52,53]. Aberrant functional organization of the Golgi is often linked to cancer. A recurrent theme in the scientific community is the hypothesis that alterations of Golgi structure are associated with cancer progression [54]. A wide range of genes induce structural alteration of the Golgi apparatus [55–57]. These alterations fall into two main categories: compaction and fragmentation of the Golgi. The available collection of genes that affect Golgi structure might form a framework for analyzing the connection between Golgi alterations and cancer.

The main challenge with genes that are linked to Golgi alterations is that they often form loosely-connected networks, making it complicated to extract meaningful information with respect to enrichment of signaling pathways or metabolic processes. To link Golgi alterations with cancer progression we used a list of 164 genes that are known to cause Golgi fragmentation (Supplementary Data S1) to filter the CTC dataset. From 164, only 97 genes were expressed. Silencing each of these gens is known to cause Golgi fragmentation [56–60]. Using the *k*-means clustering approach based on these 97 genes, cells were classified in four stable clusters (Figure 4a). Clusters 3 and 4 are highly consistent. Clusters 1 and 2, instead, have low degree of consistency.

**Figure 4.**
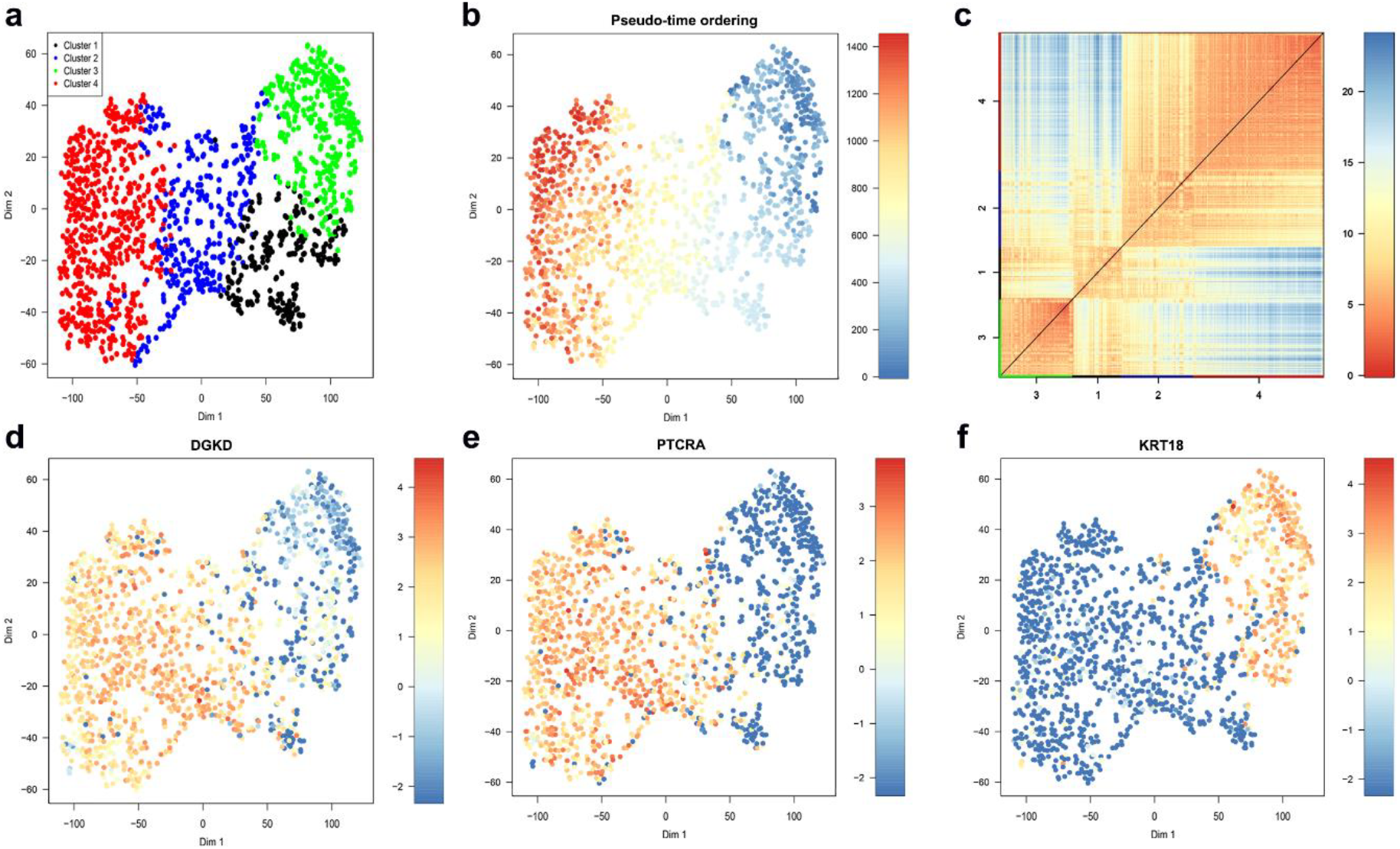
Identification of CTC sub-populations clustered based on Golgi fragmentation gene list. (a) *t*SNE map visualizing clusters of <u>circulating breast cancer cells</u> identified with *k*-means clustering. (b) The plot from (a) with cells colored based on their pseudo-time ordering. (c) A heatmap portrayal of cell-to-cell distances, cluster centers were ordered by hierarchic clustering. (d) The plot from (a) with cells colored based on their *DGKD* expression. (e) The plot from (a) with cells colored based on their *PTCRA* expression. (f) The plot from (a) with cells colored based on their *KRT18* expression.

Both pseudo-time ordering (Figure 4b) and the heatmap portrayal of cell-to-cell distances, where cluster centers were ordered by hierarchic clustering (Figure 4c), show a clear separation between clusters 3 and 4. Based on the binomial differential expression, cells in cluster 3 express significantly less *DGKD* and *PTCRA* but significantly more *KRT18* (Figure 4d-4f). Interestingly, silencing the diacylglycerol kinase, delta (*DGKD*) has been validated to cause a condensed Golgi phenotype (Chia et al, 2012). The Pre T Cell Antigen Receptor Alpha (*PTCRA*) has recently been nominated as a biomarker in breast cancer [61]. The Keratin 18 (*KRT18*) was reported as significantly less expressed in all basal-like cell lines, which are highly aggressive, compared with luminal cell lines [62]. Low *KRT18* expression in breast cancer has been suggested to correlate with poor prognosis and to be a marker for epithelial-mesenchymal transition (EMT) [63,64]. Elucidating the biological significance of these findings requires further experiments. However, we might speculate that the high *KRT18* in cluster 3 could indicate that cells in this cluster are at earlier stages of EMT and that these cells have a compact Golgi due to low *DGKD*.

The expression profiling of these 97 genes that fragment the Golgi showed two main Golgi fragmentation subsets within the Golgi genes. The first GF-subset of genes (56 genes) demonstrated up-regulation in cluster 3. The second GF-subset (15 genes) demonstrated up-regulation in cluster 4. The remaining genes (26 genes) showed either a stable low expression or unclear expression pattern across all clusters (Figure 5a). This could suggest that cells in cluster 4 exhibit a high likelihood of Golgi fragmentation whereas cells in cluster 3 exhibit a high likelihood of Golgi condensation. DEGs between clusters 3 and 4 were extracted by a two-class unpaired response test using SAMseq (FDR < 0.05 & fold change >1). Five genes (*RPS27A, TUBA1B, RPL4, RPL34* and *RPS12*) have been classified as significantly up-regulated in cluster 3 whereas 748 genes were significantly up-regulated in the cluster 4 (Supplementary Data S2). During the dissemination of breast cancer cells, they exhibit a loss of epithelial characteristics, which routinely is accompanied by up-regulation of mesenchymal genes. This process is known as epithelial-mesenchymal transition (EMT). Several studies highlighted a strong association between the expression of EMT genes in circulating breast cancer cells and cancer progression, invasion and metastasis [65,66]. To investigate this association, we profiled the expression of 20 genes (Figure 5b). These genes were nominated as EMT markers by Zhao and colleagues [67]. Seven EMT genes (*AKT1, EGFR, EPAS1, ERBB2, HIF1A, SMAD3,* and *MET*) were relatively up-regulated in cluster 3 comparing to cluster 4.

**Figure 5.**
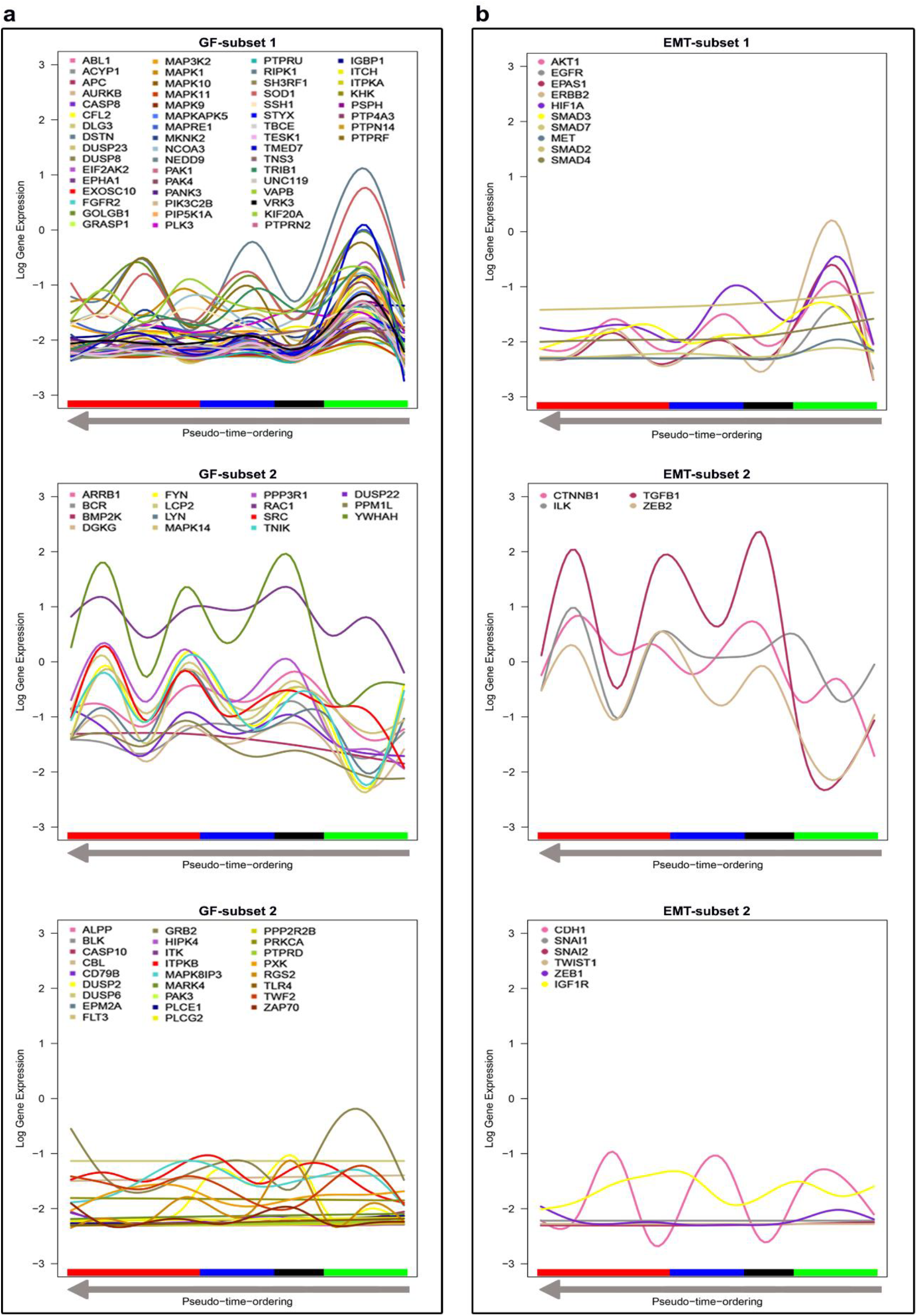
Discovery and exploration of patterns in gene expression across the clusters ordered from right to left based on pseudo-time ordering. The log gene expression values are shown on the vertical axis. Generalized additive mode smoothing is used to smooth the expression values per gene. (a) Gene expression profiling of the Golgi fragmentation gene list (97 genes). (b) Gene expression profiling of the EM gene list (20 genes).

However, four genes (*CTNNB1*, *ILK*, *TGFB1* and *ZEB2*) have been classified as significantly up-regulated in cluster 4, these genes exhibit similar expression patterns to the genes in the second GF-subset. *TGFB1* is a major inducer of EMT and is commonly used to induce EMT for research purposes [68]. *TGFB1* is highly expressed in all clusters except cluster 3. Interestingly, we found a fairly strong negative correlation (r = -0.67, p < 0.001) between *TGFB1* and *KRT18*, which is in line with previous findings [63]. Our data suggest that Golgi fragmentation is correlated with markers of EMT. On the other hand, others reported that EMT derives Golgi compaction through a strong correlation between *MMD* and *Zeb1 [69]*. However, we found weak negative correlation between the expression of *MMD* and *Zeb1* (r = -0.15, p < 0.001). This discrepancy highlights the complexity of the link between Golgi structural alterations and cancer progression and indicates the need for more experimental efforts to clarify the role of the Golgi in cancer.

### 2. MLS case study

Here, we analyse scRNA-seq data from a myxoid liposarcoma cell line. The MLS notebook is available at https://nbviewer.jupyter.org/github/ocbe-uio/DIscBIO/blob/dev/notebook/DIscBIO-MLS-Binder.ipynb. Myxoid liposarcoma (MLS) is a rare type of tumor driven by specific fusion oncogenes, normally FUS-DDIT3 [70,71], with few other genetic changes [72,73]. The 94 single cells were collected based on their cell cycle phase (G1, S or G2/M), assessed in the collection step by analyzing their DNA content using Fluorescence Activated Cell Sorter [74]. Data including ERCC spike-ins are available in the ArrayExpress database at EMBL-EBI with accession number E-MTAB-6142. During the preprocessing, genes were selected by accounting for technical noise based on the variation and expression of the ERCC spike-ins, which were added to each sample before library preparation (Figure 6a). A total of 5684 genes (black dots) have a variation above the noise level (red curve). These genes were used for the downstream analysis. After filtering the dataset, three stable MLS clusters (Jaccard similarity > 0.6) were obtained based on model-based clustering and visualized in a PCA plot (Figure 6b). Clusters can be visualized by *t*SNE maps as well. However, considering the distinctive separation, PCA plots showed intact projection of the contents of the clusters generated by model-based clustering. One possible explanation could be that the linear probabilistic processing of PCA matches with the probability model-based clustering, which generates a model for every cluster and computes the best fit of the data to the generated model [75,76]. The MLS clusters match with the cell cycle phases in the initial study [74]. Cluster one is characterized by low consistency (Silhouette width = -0.1) whereas clusters 2 and 3 have reasonable consistency (Silhouette width > 0). Pseudo-temporal ordering showed a gradual transition between the cells’ transcription profiles across the clusters (Figure 6c). Both the clustering and pseudo-time overlap with the cell cycle phases (Figure 6d). Cells in cluster 2 are mainly in G2 phase whereas the majority of cells in cluster 3 are in S phase. Cluster 1 shows heterogeneous cells with different cell cycle phases. No outlier cells were detected in any of the clusters (Figure 6e). To investigate the similarities between the single cells, Euclidean distances of Pearson transcriptome correlation matrix were computed. Based on these similarities, a heatmap portrayal of cell-to-cell distances was plotted using Euclidean as the distance measure and single linkage as the clustering method.

**Figure 6.**
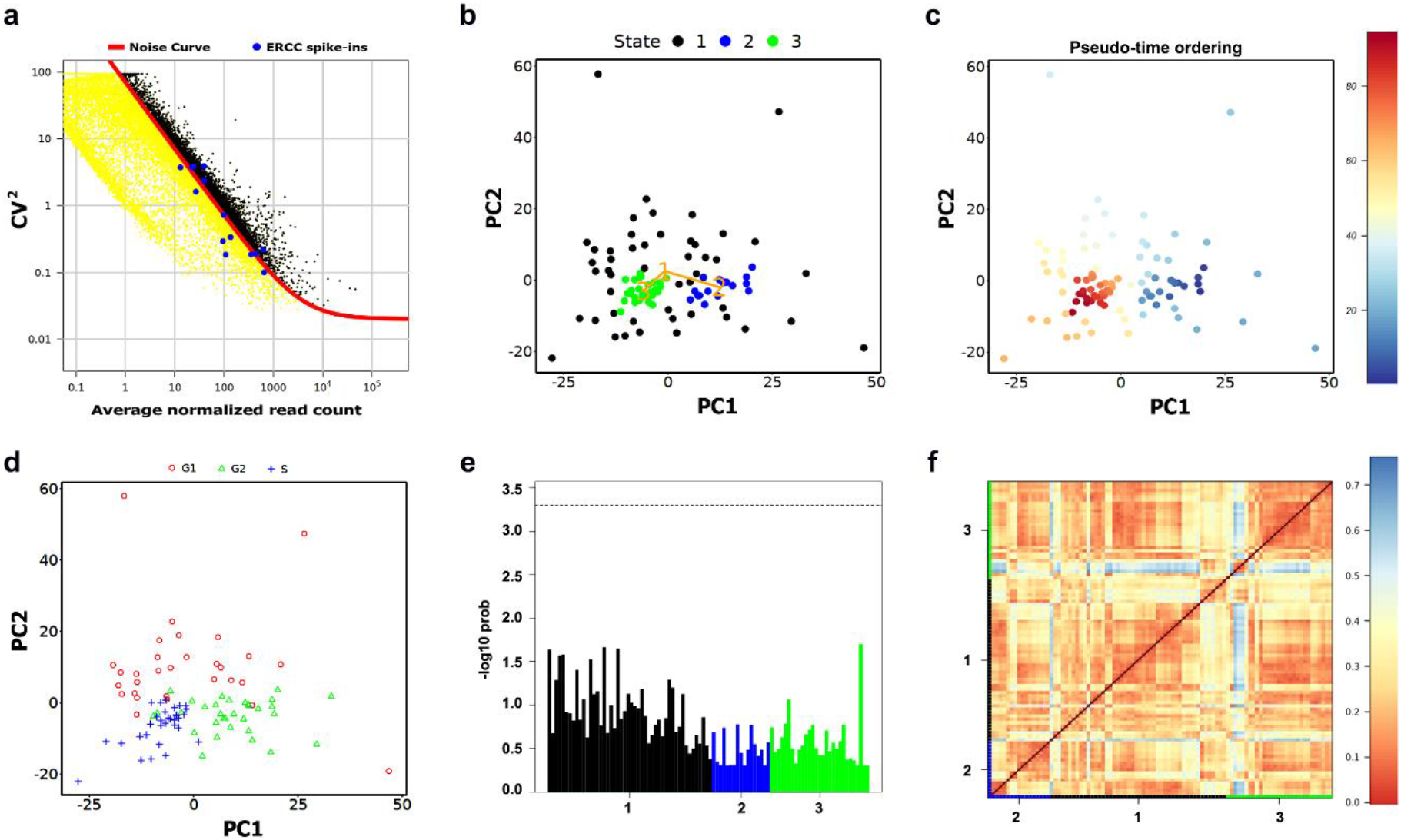
Identification of MLS sub-populations. (a) Gene filtration by accounting for technical noise based on the variation and expression of the ERCC spike-ins. (b) PCA plot visualizing clusters identified using model-based clustering. (c) The plot from (a) with cells colored based on their pseudo-time ordering. (d) The plot from (a) with cells labeled based on their cell cycle phase. (e) A bar-plot of the outlier probabilities of all cells across clusters. (f) A heatmap portrayal of cell-to-cell distances, cluster centers were ordered by hierarchic clustering.

To further define the molecular signature of the clusters, SAMseq and binomial differential expression analysis was performed (FDR < 0.05). Significant DEGs were highlighted in volcano plots (Figure 7a). Moreover, The DEGs of each cluster underwent PPI network construction. The network of the down-regulated genes in cluster 2 (Figure 7b) highlighted thymidylate synthetase (*TYMS*) as a hub gene in addition to a set of highly connected genes involved in DNA synthesis and replication. *TYMS* has been reported to be associated with the well-differentiated subtype of liposarcoma [77]. *TYMS* is up-regulated in cluster 3 (fig 7c). To identify potential genes to predict the cluster identity of a target cell whether it belongs to cluster 2 or 3, decision tree analysis was implemented. The generated RPART decision tree (fig 7d) included two decision nodes: Aurora Kinase A (*AURKA*) and Cell Division Cycle 27 (*CDC27*). AURKA is involved in self-renewal of breast cancer stem cells [78]. *CDC27* is known to induce metastasis, invasion and sphere-formation in colorectal and gastric tumors [79,80]. Cluster 2 cells, which were ordered in the foreground pseudo-temporal ordering, highly express genes involved in regulating proliferation, EMT, stemness and chemoresistance acquisition in several cancers. Stemness is known to be frequently associated with quiescence [81]. Several studies reported that the decision for entering quiescence is facilitated during the maternal G2 phase [82,83]. G2 arrest in stem cells is associated with robust regeneration capacity and it has been reported to be reversible and released when regeneration must be achieved [84]. The networking analysis outcome (Supplementary Table. S1) nominated a panel of 18 hub genes, including *PLK1*, *CDC20*, *KIF2C*, *BUB1*, *AURKB*, and *PTTG1* as potential biomarkers for cluster 2 cells. The overall impact of our findings reported here is that cells in cluster 2, which are mainly in G2 phase, seem to exhibit aggressive and stem-like properties and further computational and experimental investigations should validate these biomarkers candidates in myxoid liposarcoma.

**Figure 7.**
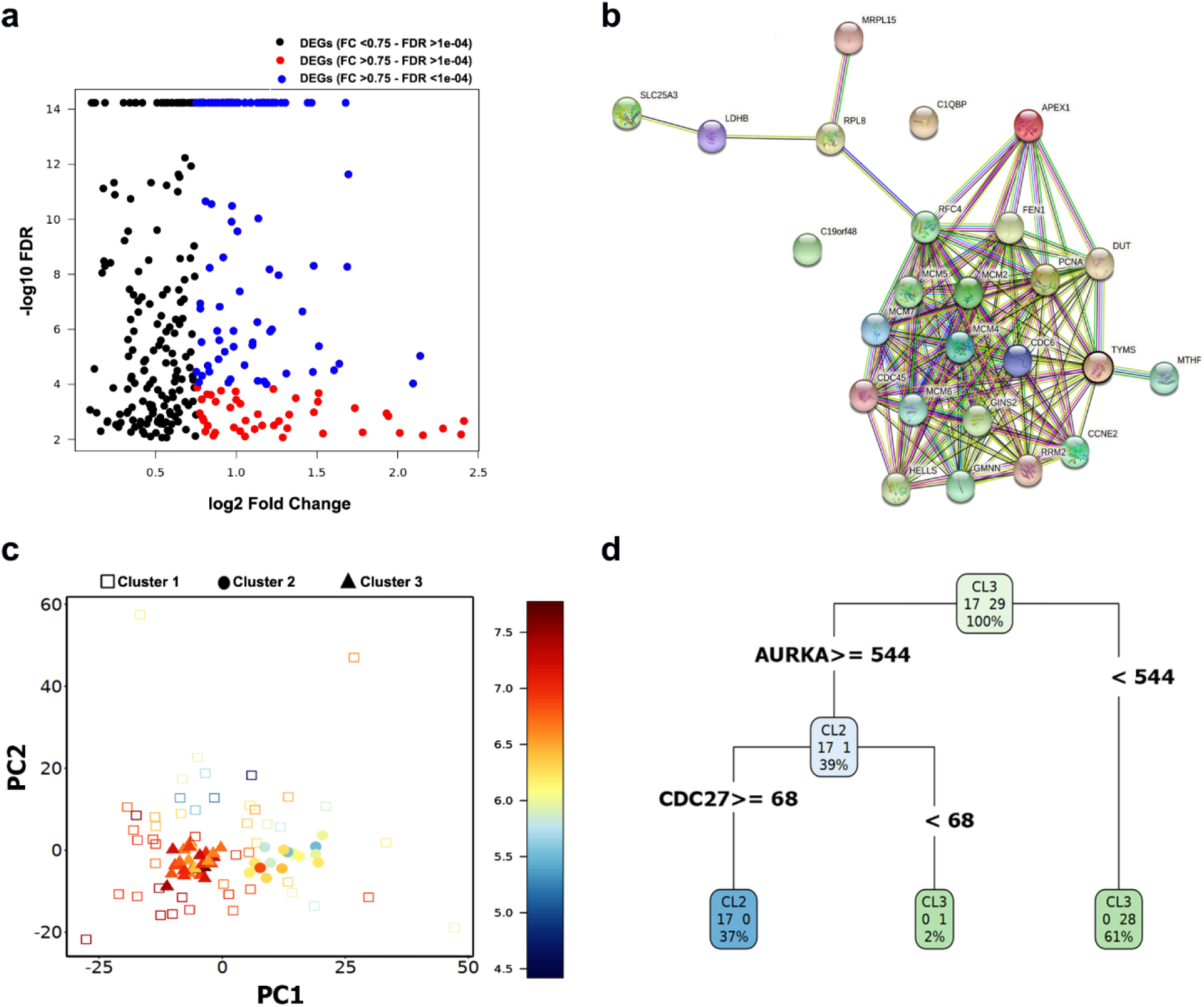
Biomarker discovery in cluster 2. (a) Volcano plot showing the up-regulated markers in cluster 2. These markers were detected using binomial differential expression. (b) Network of down-regulated DEGs in cluster 2. (c) PCA plot visualizing cells labeled based on their cluster ID and color-coded based on the expression of *TYMS*. (d) Schematic figure explaining the RPART decision tree to predict the cluster identity of a target cell to belong to cluster 2 or 3.

## Conclusion

To reduce the complexity of single-cell transcriptomics analyses using a combination of specific computational tools for scRNA-seq analysis, we developed DIscBIO as a step-wise approach. It is available as an R package, Jupyter notebook and Binder cloud version. This makes it a convenient tool for a variety of researches to facilitate the exploration of their scRNA-seq data. Furthermore, and due to its sequential narrative design and explanatory structure, DIscBIO can be used for teaching and training purposes. Using DIscBIO we were able to identify CTCs with highly aggressive characteristics of invasion and metastasis in breast cancer and also gain insights about the link between the alterations of Golgi apparatus and cancer progression. Furthermore, DIscBIO enabled us to identify a small subset of cells with possible aggressive and stem-like properties in myxoid liposarcoma.

## Data availability

DIscBIO is an open-source pipeline whose source code and Jupyter notebooks are deposited in a GitHub repository https://github.com/ocbe-uio/DIscBIO. Its CRAN-published R package can be downloaded from https://cran.r-project.org/web/packages/DIscBIO/index.html or installed directly from a regular R session. DIscBIO is also provided as a cloud version using Binder. Instructions to run our Binder notebooks can be found in the README on our GitHub repository.

## Supplementary Materials

The following are available online at www.mdpi.com/xxx/s1, Supplementary Table S1: Networking analysis outcome of DEGs in cluster 2 of myxoid liposarcoma cells, Supplementary Data S1: a list of 164 genes that are known to cause Golgi fragmentation, Supplementary Data S2: GEDs in CTC dataset.

## Author Contributions

Conceptualization, A.K.L., A.S., H.F., and. S.G.; software, S.G., W.L.N., B.R.K. and D.F.; methodology, H.F., E.J., S.G., A.P. and A.K.L.; analysis, S.G.; investigation, W.L.N., S.G., and A.P.; visualization, S.G., B.R.K. and A.K.L.; writing—original draft preparation, S.G., A.K.L., and H.F.; writing—review and editing, S.G., D.F., W.L.N., E.J., A.S., H.F. and A.K.L.; supervision, A.K.L., H.F. and A.S. All authors have read and agreed to the published version of the manuscript.

## Acknowledgement

We would like to thank Dominic Grün, Benjamin Ulfenborg and Hongkai Ji for their kind feedback and input during the construction of this pipeline. A.K.L. acknowledges the Centre for Digital Life Norway for supporting the partner projects PerCaThe and BigInsight.

