## Supplementary Table S1 for "DIscBIO: a user-friendly pipeline for biomarker discovery in single-cell transcriptomics"

### DIscBIO: a user-friendly pipeline for biomarker discovery in single-cell transcriptomics

**Salim Ghannoum[1*], Waldir Leoncio Netto[2], Damiano Fantini[3], Benjamin Ragan-Kelley[4], Amirabbas Parizadeh[1], Emma Jonasson[5], Anders Ståhlberg[5, 6, 7], Hesso Farhan[1] and Alvaro Köhn-Luque[2*]**

1 Department of Molecular Medicine, Institute of Basic Medical Sciences, University of Oslo, Norway.

2 Oslo Centre for Biostatistics and Epidemiology, Faculty of Medicine, University of Oslo, Norway.

3 Department of Urology, Northwestern University, Chicago, USA.

4 Simula Research Laboratory, Lysaker, Norway.

5 Sahlgrenska Center of Cancer Research, Department of Laboratory Medicine, Institute of Biomedicine, Sahlgrenska Academy at University of Gothenburg, Gothenburg, Sweden.

6 Wallenberg Centre for Molecular and Translational Medicine, University of Gothenburg, Sweden.

7 Department of Clinical Genetics and Genomics, Sahlgrenska University Hospital, Gothenburg, Sweden.

**Table. S1:** Networking analysis outcome of DEGs in cluster 2 of myxoid liposarcoma cells showing the genes high in both connectivity degree and betweenness centrality in addition to their known functions.

| Genes | Connectivity degree | Betweenness centrality | Known Functions | References |
|---|---|---|---|---|
| **PLK1** | 80 | 100.64 | ❖ Stemness biomarker candidate in bladder cancer.<br>❖ Affecting the chemoresistance of ovarian cancer. | [1]<br>[2] |
| **CDC20** | 79 | 96.76 | ❖ A master cell cycle regulator<br>❖ Maintaining the self-renewal ability of prostate cancer stem cells. | [3]<br>[4] |
| **CDK1** | 77 | 67.07 | ❖ A regulator of tumor-initiating in melanoma. | [5] |
| **CCNB1** | 76 | 22.19 | ❖ Aactivate EMT. | [6] |
| **KIF2C** | 76 | 76.55 | ❖ Mediating chemoresistance in ovarian cancer cells. | [7] |
| **KIF11** | 75 | 28.18 | ❖ Critical for self-renewal and stemness in breast cancer. | [8] |
| **CENPE** | 75 | 16.79 | ❖ Promotes sphere forming ability in glioblastoma. | [9] |
| **MAD2L1** | 75 | 28.24 | ❖ Modulates pluripotency and stemness ability in Gastric cancer. | [10] |
| **CCNB2** | 74 | 25.51 | ❖ Associated with cancer stem cell properties in lung squamous cell carcinoma. | [11] |
| **BUB1** | 74 | 56.97 | ❖ Maintaining cancer stem cells in breast cancer.<br>❖ Promoting proliferation and radio-resistance in glioblastoma. | [12]<br>[13] |
| **AURKA** | 73 | 3.49 | ❖ Involved in self-renewal of breast cancer stem cells. | [14] |
| **BUB1B** | 73 | 23.07 | ❖ Promotes tumor proliferation and induces radioresistance in glioblastoma. | [15] |
| **AURKB** | 73 | 39.61 | ❖ Stemness biomarker candidate in bladder cancer. | [1] |
| **CDCA8** | 73 | 52.94 | ❖ Promoting melanoma malignant progression. | [16] |
| **NUF2** | 72 | 17.88 | ❖ Closely linked with the progression of pancreatic cancer. | [17] |
| **CENPF** | 72 | 21.95 | ❖ Promoting the aggressiveness of gastric cancer cells. | [18] |
| **CCNA2** | 71 | 75.37 | ❖ A growth regulator in colorectal cancer. | [19] |
| **TPX2** | 71 | 66.90 | ❖ Closely related to the development of breast cancer stem cells. | [20] |
